# Nucleoside-driven specificity of DNA Methyltransferase

**DOI:** 10.1101/2022.10.26.513954

**Authors:** Madhuri Gade, Jasmine M. Gardner, Prashant Jain, Paola Laurino

## Abstract

We have studied adenosine binding specificities of two bacterial DNA methyltransferase Taq methyltransferase (M.TaqI), and HhaI methyltransferase (M.HhaI). While these DNA methyltransferases have similar cofactor binding pocket interactions, experimental data showed different specificity for novel cofactors ((SNM) (S-guanosyl-L-methionine (SGM), S-cytidyl-L-methionine (SCM), S-uridyl-L-methionine (SUM)). Protein dynamics corroborate the experimental data on the cofactor specificities. For M.TaqI the specificity for S-adenosyl-L-methionine (SAM) is governed by the tight binding on the nucleoside part of the cofactor, while for M.HhaI the degree of freedom of the nucleoside chain allows the acceptance of other bases. The experimental data proves a catalytically productive methylation by M.HhaI binding pocket for all the SNM. Our results suggest a new route for successful design of unnatural SNM analogues for methyltransferases as a tool for cofactor engineering.

**Table of Content:** **TOC**: Methylation by DNA methyltransferase is nucleobase dependent. While M.Taq1 is specific for SAM, M.HhaI is promiscuous for other SAM analogs

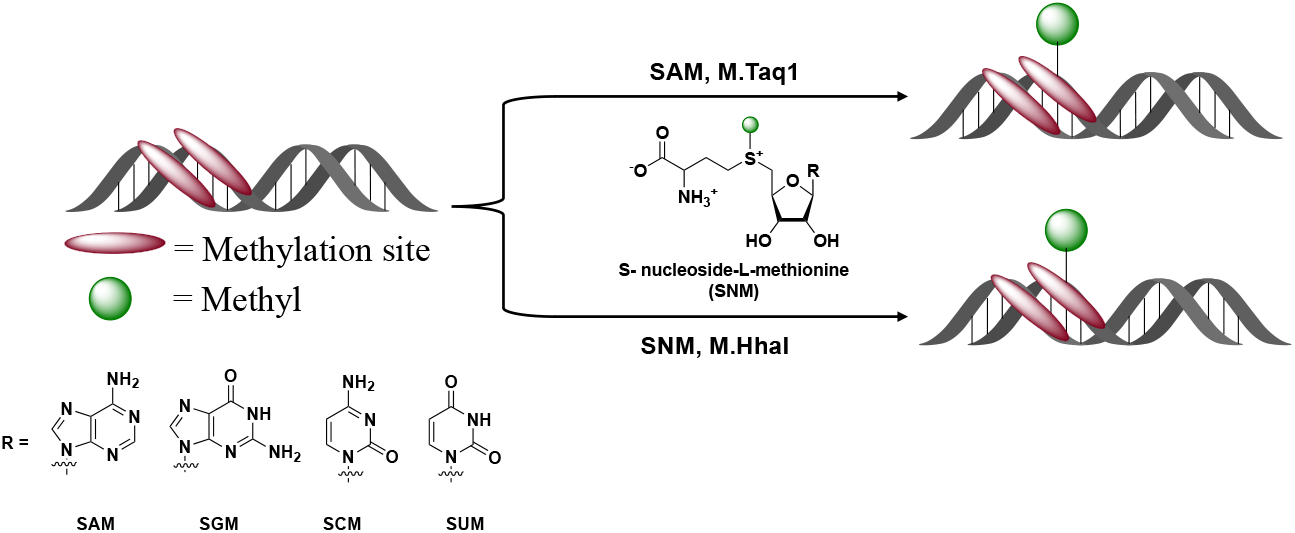

## Introduction

Nucleotides and their derivatives are the most common chemical moiety present in natural ligands^[1]^. Adenosine moiety is the most abundant among the nucleotides and plays a key role in all the living organisms^[2–3]^. Adenosine’s role as a cofactor or ligand in various cellular processes is fulfilled by promoting specific interactions with proteins and enzymes. These specific interactions allow proteins to distinguish between adenosine and other ligands or nucleotides^[2, 4–5]^. Ultimately adenosine specific interactions are exploited by the cell for the fine regulation of enzymes in the context of metabolic processes^[4]^. Adenosine moiety is overrepresented in cofactors, namely adenosine 5′-triphosphate (ATP), nicotinamide adenine dinucleotide (NAD), flavin adenine dinucleotide (FAD), S-adenosyl methionine (SAM), and coenzyme A (CoA). Although SAM is involved in various biological processes in cells^[6–9]^, its main role is methylation of biomolecules^[10]^. This methylation is facilitated by a class of enzymes called methyltransferases^[11]^.

Methyltransferase is a large family of enzymes that catalyzes the transfer of a methyl group from SAM to biomolecules (DNA, RNA, protein, and small molecules)^[12–13]^ (Figure 1a). Methylation of these biomolecules plays a critical role in various processes like gene expression and regulation, signal transduction, gene silencing, and chromatin regulation^[14–15]^. One important class of methyltransferases is DNA methyltransferase^[16–17]^. For instance, in prokaryotes, DNA methyltransferase has a crucial role in distinguishing between endogenous and exogenous DNA^[18]^. In mammals, DNA methylation plays a key role in embryonic development and gene regulation^[19]^. Considering the significance of DNA methylation, it is essential to find an easy method to detect methylation patterns in cells.

**Figure 1.**
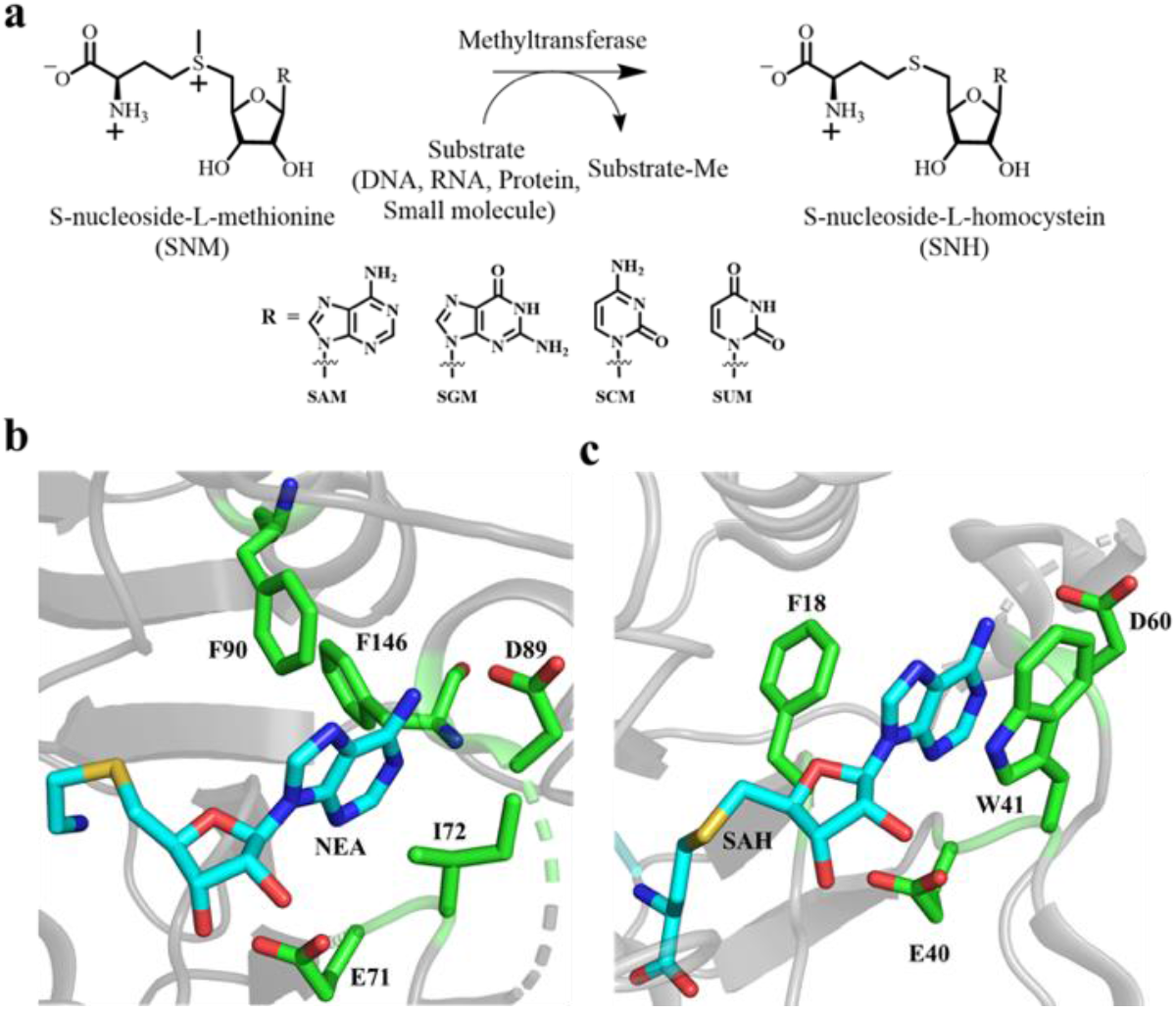
Natural methylation reaction and binding pockets of M.TaqI and M.HhaI DNA methyltransferase. (a) Schematics of transfer of methyl group from cofactor to biomolecules (DNA, RNA, protein, and small molecules) and structures of S-nucleoside methionine (SNM) cofactor used in this study. (b) Binding site residues of M.TaqI interacting with adenosine of 5’-Deoxy-5’-[2-(amino)ethylthio]adenosine (NEA). NEA is in cyan and interacting residues are in green color (PDB ID 1g38). (C) Binding site residues of M.HhaI interacting with adenosine of SAH. SAH is in cyan and interacting residues are in green (PDB ID 2hr1).

One approach includes the use of SAM analogues that carry an activated transferring group which will allow the labeling of the substrates in cells and ultimately the study of the methyltransferase’s function^[20–24]^. However, use of SAM analogs remains limited because of the lack of biorthogonality ^[25–26]^ for instance the specific recognition of synthetic cofactor that ultimately allows fine regulation of an engineered methyltransferase.

As mentioned above, adenosine shows specific interactions with protein. In DNA methyltransferases, adenosine moiety has several sites for hydrogen bonding (acceptors N1, N3, and donor N6 of adenine and ribose hydroxyl group) and pi-pi stacking interaction with adenine aromatic ring^[2]^. N6 of adenine is primarily involved in hydrogen bonding with the side chain of aspartate residue^[3]^. It is well documented that ribose hydroxyl forms bidentate interaction with aspartate or glutamate^[27]^. Indeed, interactions between SAM and DNA methyltransferases at the active site are essential for the specificity of the enzyme. By taking advantage of these specific interactions, one could think of replacing the adenosine moiety, achieve specificity and ultimately orthogonality for a SAM analogue carrying a different nucleobase. A recent report functionalized adenosine with different lateral group, however no changes of the aromatic core is reported so far^[20]^. At the same time a detailed understanding of interactions is required to analyze factors governing cofactor discrimination. Recognition between cognate and non-cognate SAM analogs by the DNA methyltransferases is usually elucidated by sequence specificity that allows specific protein-cofactor interactions and steric hindrance in the cofactor binding pocket^[28]^. Still, it is less understood if enzyme dynamics play a role in differentiating between cofactor analogs.

Herein, we have investigated the adenosine binding site for two DNA methyltransferases by utilizing novel S-nucleoside methionine (SNM) (S-guanosyl-L-methionine (SGM), S-cytidyl-L-methionine (SCM), S-uridyl-L-methionine (SUM)) cofactors^[29]^. These SNM cofactors carry different nucleotide bases, making them ideal candidates to study the adenosine binding pocket of DNA methyltransferase. This study opens a new avenue for the design of SAM derivatives carrying different nucleobases and their potential to achieve biorthogonality because of substantial changes in the cofactor structure.

## Results and Discussions

We tested two representative classes of DNA methyltransferase Taq methyltransferase (M.TaqI), and HhaI methyltransferase (M.HhaI). M.TaqI recognize the DNA sequence TCG**A** methylating N6 of the adenine base^[30]^, while M.HhaI recognizes G**C**GC sequence methylating the C5 cytosine of the cytosine base ^[31]^. In M.TaqI, adenosine N1 and N6 interacts via hydrogen bonding with F90, D89 respectively. Adenine ring forms Van der Waals interaction with the side chain of I72, F90, and F146. The ribose hydroxyl groups have a bidentate interaction with E71^[32]^ (Figure 1b). In the case of M.HhaI adenosine, N6 forms hydrogen bonding with D60. Adenine ring of SAH has pi-pi staking interaction with W41 (parallel) and F18 (perpendicular) (Figure 1b). The ribose hydroxyl groups have bidentate interaction with E40^[33]^. M.TaqI and M.HhaI have identical key interactions with the adenosine group of SAM, this makes them optimal candidates to investigate the contribution of the enzyme dynamics for cognate and non-cognate cofactor preference.

To investigate the specificity of M.TaqI and M.HhaI for different SNM (SAM, SGM, SCM and SUM) analogs, we utilized the enzymatically synthesized SNM cofactors by hMAT2A^[29]^. To test the methylation activity of M.TaqI and M.HhaI for the different SNM, we used DNA protection against the restriction enzyme assay^[34]^ (Figure 2a). Modification of DNA with methyl group by methyltransferase prevents fragmentation by corresponding restriction enzymes that are methylation-sensitive^[35]^. First, we set up the reaction of hMAT2A with NTPs (ATP, GTP, CTP, and UTP) and methionine for one hour then, hMAT2A was inactivated, followed by the addition of DNA substrate and DNA methyltransferase. We used dsDNA substrate for M.TaqI and M.HhaI, with one recognition site for methylation. Using dsDNA substrate and various cofactor (SAM, SGM, SCM, and SUM), we carried out a methylation reaction using M.TaqI at 65 °C and M.HhaI at 37 °C for one hour followed by digestion with restriction enzymes TaqI and HhaI respectively for one hour and run on the agarose gel. This assay demonstrates that M.TaqI is specific for SAM, though 50% activity was observed for SCM too (Figure 2b). Whereas DNA treated with SGM, and SUM was susceptible for TaqI digestion, and two bands were detected on the gel (Figure 2b). To exclude that the observed specificity is due to SNM instability, we tested the degradation of SNM analogs at 65 °C over time. If the degradation is fast, SNM would be unavailable for the reaction. SNM degrade similarly and up to 90% over 60 minutes at 65 °C. However, in the current reaction condition still 100 μM of SNM were available for the reaction compared to the initial concentration (Figure S1). The K_M_ of SAM for M.Taq is 3.7 μM^[36]^, the amount left in the reaction mixture is well above the required for the methylation reaction.

**Figure 2.**
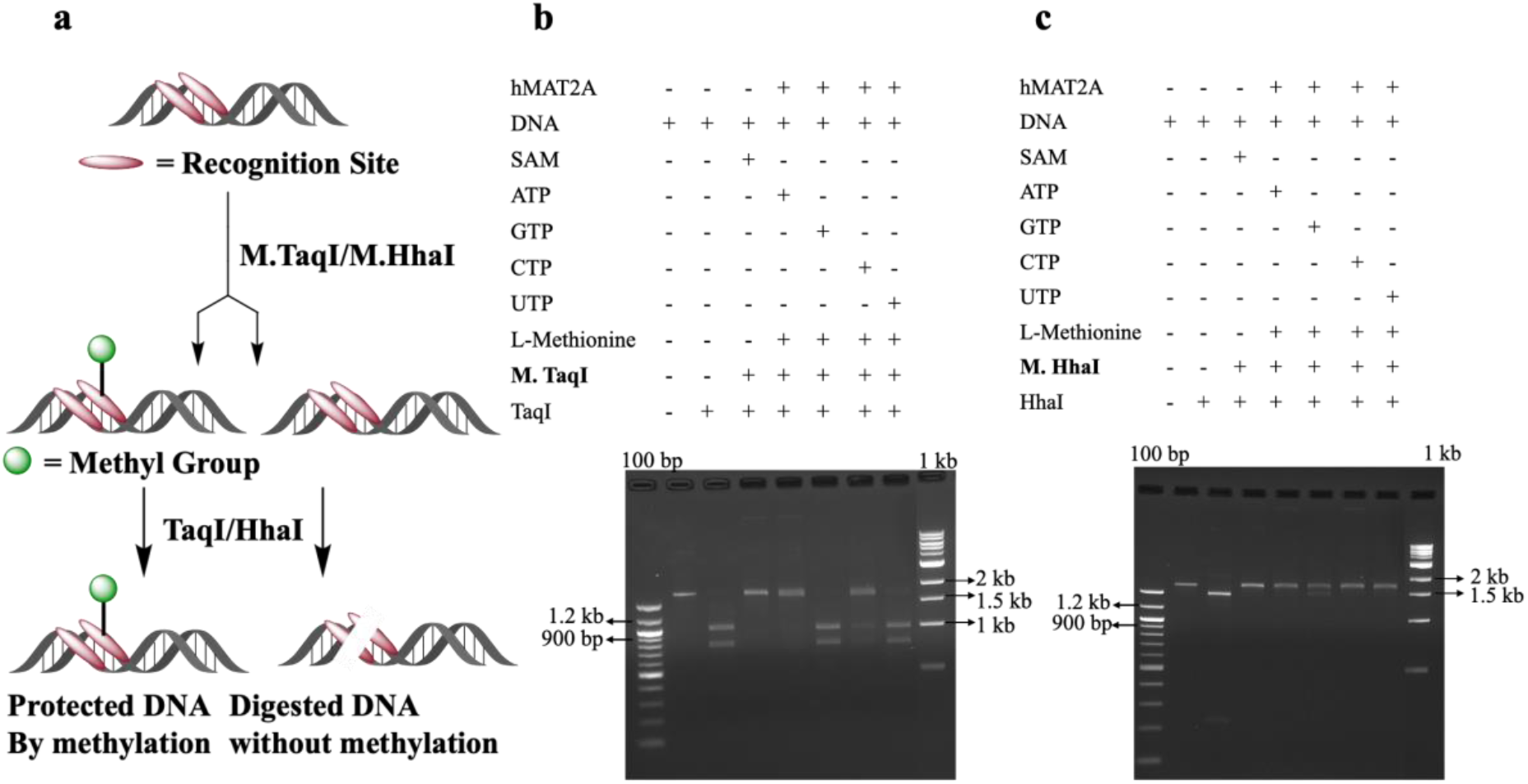
Agarose gel analysis of DNA modification. Enzymatic modification of (a) M.TaqI and (b) M.HhaI byNM cofactors. Agarose gel was 1.5 % run for 1 hr. at 100 V, stained with ethidium bromide. GeneRuler DNA ladder 100bp and 1kb was used.

M.HhaI was observed to accept SGM, SCM, and SUM cofactors for the methylation reaction. dsDNA substrate was protected from digestion with the corresponding restriction enzyme HhaI in the case of SGM, SCM and SUM (Figure 2c). Indeed, the gel showed single band as outcome of the digestion reaction (Figure 2c). The relative efficiency of protection is similar for four different SNM cofactors for M.HhaI by gel. No spontaneous methylation without M.TaqI and M.HhaI was observed using different SNM analogs in these conditions (Figure S2).

Further, we investigated the molecular basis for observed specificity for M.TaqI and promiscuity for M.HhaI by molecular dynamic (MD) simulation. For each representative structure, an all-atom explicit water system was created starting from the relevant crystal structure (PDB IDs 1g38 and 1skm for M.TaqI and M.HhaI, respectively). All ligands, cofactors, and DNA were removed to produce the apo protein structure. To produce the bound structures SNH co-factors were added in alignment with the original bound cofactor. The protein was solvated in a cubic water box padded with 10Å on each side and neutralized with Cl ions. MD simulations were performed using Gromacs 2020v.3^[37]^. The Amber 14 forcefield^[38]^ was used with TIP3P water^[39]^. A modified Berendsen thermostat^[40]^ maintained the temperature at 300K and a Parrinello-Rahman barostat^[41]^ maintained a pressure of 1 bar. A 2 fs timestep was employed. Each system was equilibrated for 50 ps in NVT environment followed by 50 ps in NPT environment. The first 10 ns of simulation time were discarded for equilibration purposes. The systems were simulated for an additional 500 ns of production simulation time in an NPT environment with 5 independent replicates. SNH cofactors stayed bound for a minimum of 10 ns indicating a stable bound starting structure. Longer time-scale binding varied between proteins and cofactors. To observe differences in binding patterns, simulations are aligned around the binding pocket defined as alpha carbons of residues within 7 Å of the crystal structure bound SAH. RMSF of ligand heavy atoms is calculated on the SNH analogs in relation to the aligned structures.

Cofactor binding patterns for M.HhaI and M.TaqI can be seen in Figures 3 and 4 as observed by hydrogen bonding patterns. In M.HhaI and M.TaqI, the most highly conserved binding mode is the conserved bidentate interaction between the SNH-analog ribose moiety and a glutamate residue (Glu-40 and Glu-71 in M.HhaI and M.TaqI, respectively). Despite strong conserved binding with the ribose moiety in each observed SNH-analog, the binding pattern is not sufficient to maintain cofactor binding, as observed by SUH bound to M.TaqI. In M.TaqI, SUH binds strongly between the ribose moiety and Glu-71, but without additional binding modes in other cofactor moieties, binding is not maintained for more than 50 ns in the MD simulations. Other than the conserved bidentate interaction, M.HhaI and M.TaqI exhibit very different binding modes.

**Figure 3.**
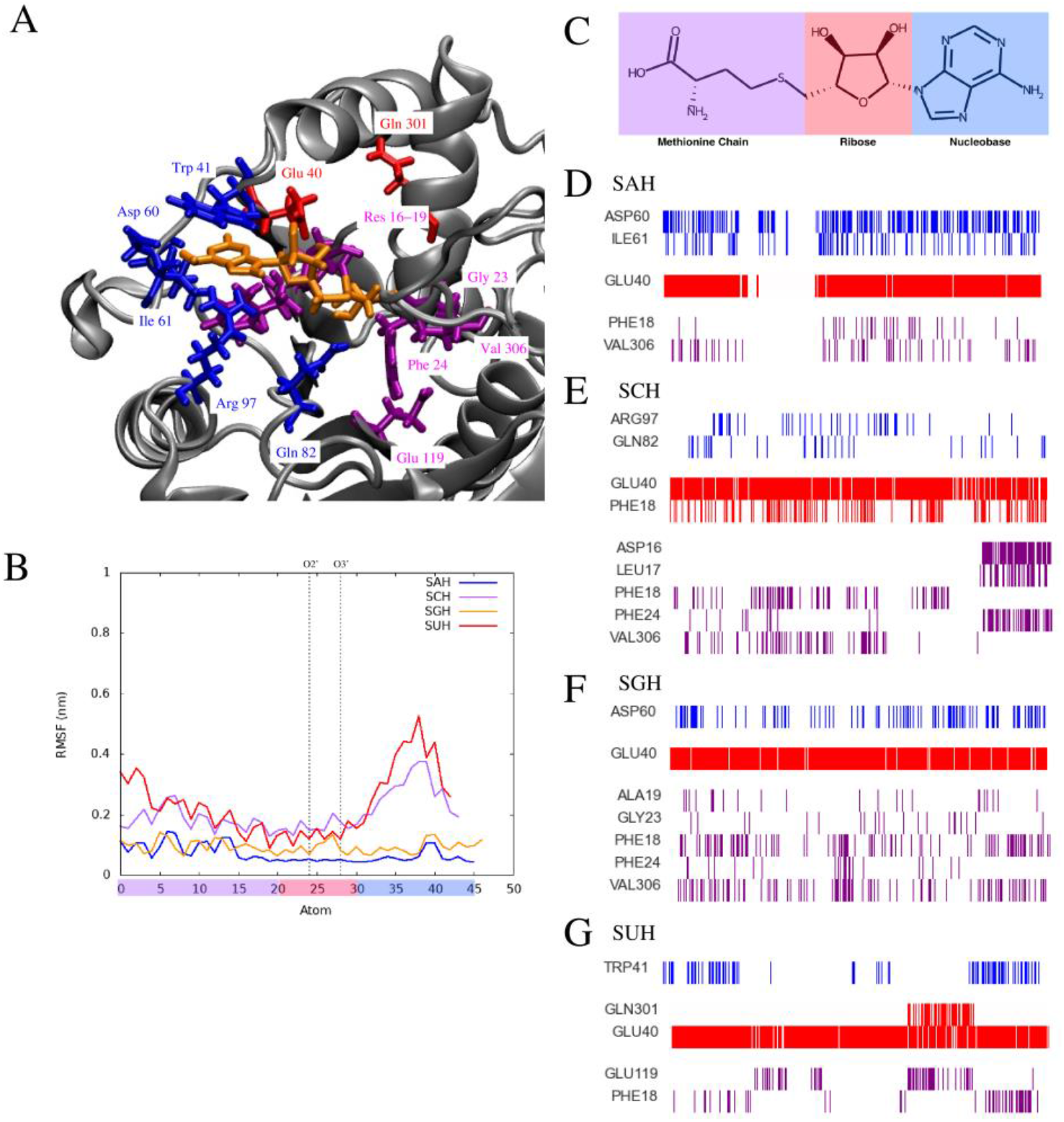
Hydrogen bonding patterns of M.HhaI bound with SNH-analogs. In each image, the atoms are coloured corresponding to their categorisation within the SAH molecule where purple indicates the methionine chain, red indicates ribose, and blue indicates nucleobase atoms. Only residues with sustained binding, or binding for more than 10% of the simulation time, are shown. A: Representative snapshot of SAH bound to M.HhaI with binding residues shown in liquorice representation and coloured corresponding to binding locale: blue for nucleobase, red for ribose, and purple for methionine chain. SAH molecule is shown in orange. B: RMSF of heavy atoms on analog SNH cofactors averaged over a single 500 ns simulation. SAH is shown in blue, SCH in purple, SGH in orange, and SUH in red. Atom numbering begins at the methionine chain and moves progressively to the nucleobase. The number scale is coloured according to the atom categorisation into methionine chain (purple), ribose (red), or nucleobase (blue). Atoms which participate in the conserved bidentate interactions are denoted by dashed lines on the plots. C: Representative SAH molecule with atom type highlighted corresponding to its categorisation in figures D-G. D-G: Binding existence over 5 independent simulations of 500 ns each for a total of 2,500 ns of simulation time. Each vertical line represents a snapshot after 1 ns of simulation time. Plots are shown for SAH (D), SCH (E), SGH (F), and SUH(G).

**Figure 4.**
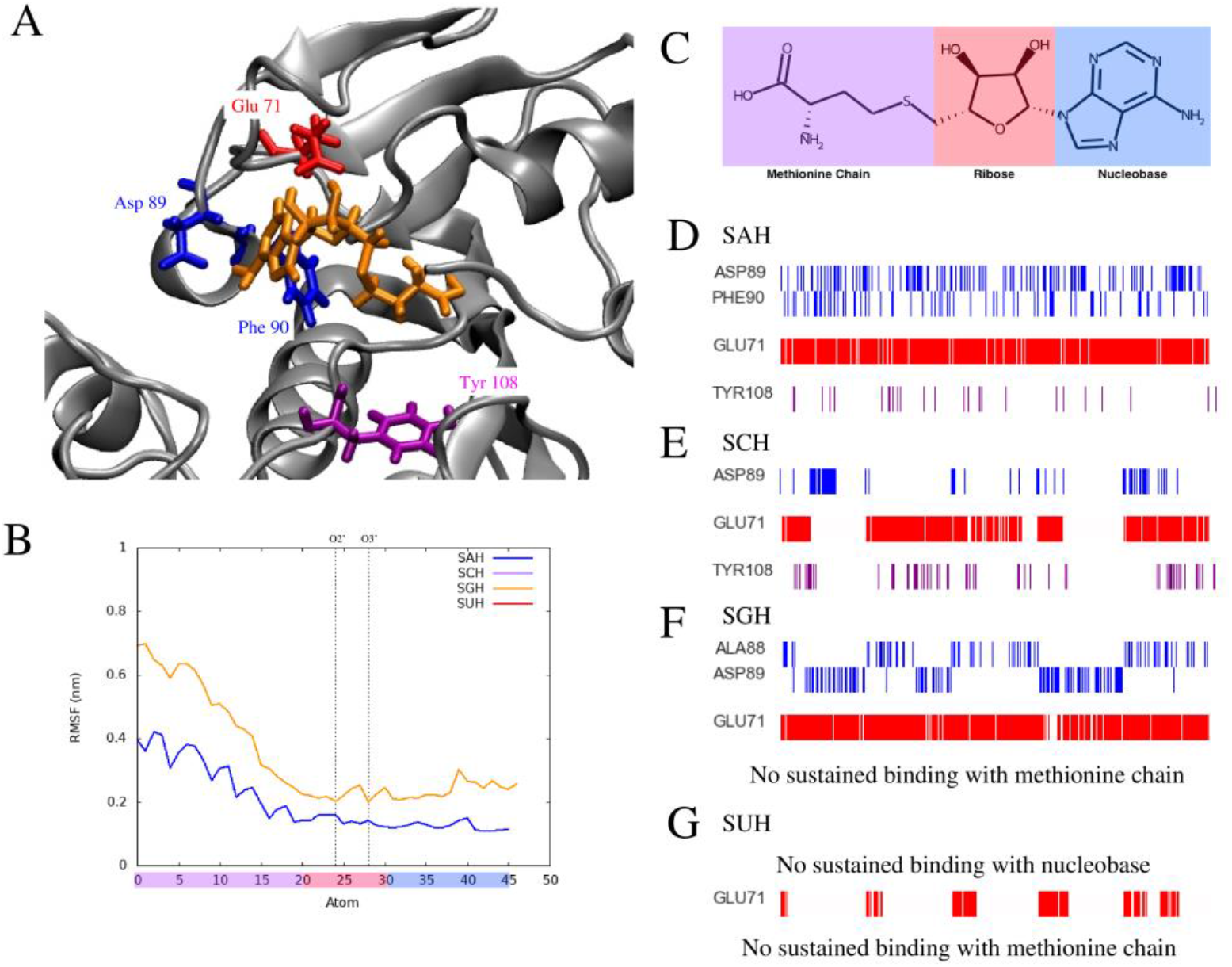
Hydrogen bonding patterns of M.TaqI bound with SNH-analogs. In each image, the atoms are coloured corresponding to their categorisation within the SAH molecule where purple indicates the methionine chain, red indicates ribose, and blue indicates nucleobase atoms. Only residues with sustained binding, or binding for more than 10% of the simulation time, are shown. A: Representative snapshot of SAH bound to M.TaqI with binding residues shown in liquorice representation and coloured corresponding to binding locale: blue for nucleobase, red for ribose, and purple for methionine chain. SAH molecule is shown in orange. B: RMSF of heavy atoms on analog SNH cofactors averaged over a single 500 ns simulation. SAH is shown in blue and SGH is shown in orange. SCH and SUH are not shown due to unbinding events in each run. Atom numbering begins at the methionine chain and moves progressively to the nucleobase. The number scale is coloured according to the atom categorisation into methionine chain (purple), ribose (red), or nucleobase (blue). Atoms which participate in the conserved bidentate interactions are denoted by dashed lines on the plots. C: Representative SAH molecule with atom type highlighted corresponding to its categorisation in figures D-G. D-G: Binding existence over 5 independent simulations of 500 ns each for a total of 2,500 ns of simulation time. Each vertical line represents a snapshot after 1 ns of simulation time. Plots are shown for SAH (D), SCH (E), SGH (F), and SUH(G).

In M.HhaI, binding occurs throughout the SNH-analog, including numerous hydrogen bonding partners available to the methionine moiety (Figure 3). SAH binds the nucleotide tightly with the sidechain of Asp-60 and alternative binding with the backbone of Ile-61, while SGH shows looser binding only to Asp-60. SUH and SCH indicate more sparse binding in the nucleotide moiety, with SUH binding to the backbone of TRP-41 and SCH binding to the sidechains of Arg-87 and Gln-82. All analogs indicate varied yet sustained binding of the methionine moiety. The RMSF of the SNH-analogs in the binding pockets (Figure 3B) show a clear lack of flexibility of the methionine moiety in binding patterns with higher flexibility of the nucleotide moiety in SCH and SUH. The most highly conserved interaction undergone by the methionine moiety is hydrogen bonding with the backbone of Phe-18 with this binding pattern observed in each analog. Additional hydrogen bonding of the methionine moiety occurs through backbone atoms of residues 16-19 and 23-24. Backbone hydrogen bonding is also observed with Val-306 in SAH, SCH, and SGH, while Glu-119 interacts with SUH through sidechain interactions.

M.TaqI exhibits very different binding patterns as seen in Figure 4. In binding with M.TaqI, SAH binds strongly through the nucleobase moiety, while SGH indicates moderate nucleobase binding. SCH and SUH indicate very poor binding in the nucleobase moiety and further in the molecule. The M.TaqI binding pocket provides high flexibility to the methionine chain, leaving the burden of binding to the nucleobase and ribose moieties and little to no hydrogen bonding activity for the methionine moiety. The only observed binding with the methionine moiety is a sparse backbone interaction with Tyr-108 observed in SAH and SCH. The nucleobase moiety hydrogen bonds with the sidechain of Asp-89, a residue analogous to Asp-60 in M.HhaI with sustained interactions observed in binding with SAH and SGH and looser interactions observed with SUH. The nucleobase moiety on SAH participates in alternative binding with the backbone of Phe-90 while SGH sparsely binds with the backbone of Ala-88 when not bound to Asp-89.

The binding patterns are further exemplified in the length of binding for each SNH analog. Binding patterns may be qualitatively observed in Figure 3 and 4 as gaps in hydrogen bonding throughout all moieties. In M.HhaI, each of the SNH analogs maintained bound conformations throughout the 500 ns simulations, with the exception of a single replica of SAH binding. This unbinding event is not expected to be indicative of physical phenomena, but rather a result of normal fluctuations and relatively rare events which can happen due to the stochasticity of MD simulations. In M.TaqI, SAH and SGH maintained a bound position throughout the 500 ns simulations while SUH and SCH exhibited very poor binding. SUH unbound from its initial position within 50 ns in each replica. SCH remained bound in only 2 out of the 5 replicas for the complete 500 ns.

Though binding is conserved through similar bonding interactions in M.TaqI with SAH and SGH analogs, enzymatic activity is not observed with SGM in physical experiments. Protein fluctuations were studied using Principal Component Analysis (PCA) to reduce dimensionality of the conformational space. Dimensionality reduction was performed on protein alpha carbon atoms using 2,500 total ns of simulation time with the Gromacs 2020v3 software package^[37]^. The resulting populations of protein conformations over the trajectory in the first and second principal components (PC 1 and PC 2) are shown in Figure 5 and 6).

**Figure 5.**
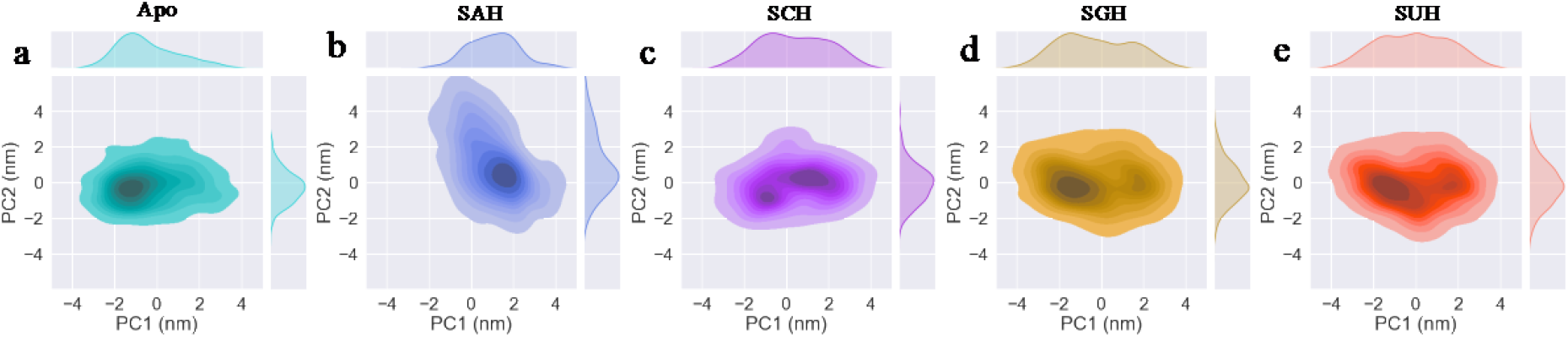
Conformations of 2,500 ns of simulation time plotted on the first and second principal components for M.TaqI in the apo form (a) and bound with SAH (b), SCH (c), SGH (d), and SUH (e). Concentration of color indicates density.

**Figure 6:**
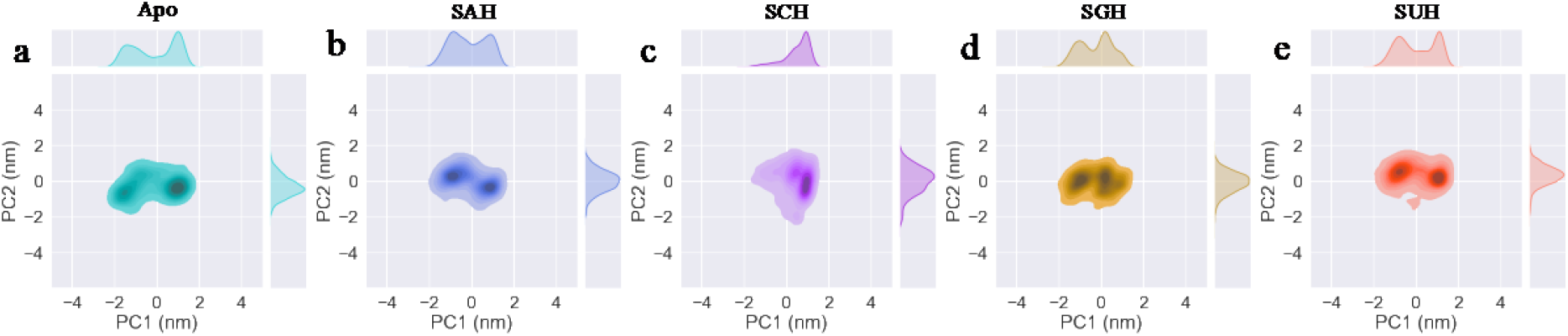
Conformations of 2,500 ns of simulation time plotted on the first and second principal components for M.HhaI in the apo form (a) and bound with SAH (b), SCH (c), SGH (d), and SUH (e). Concentration of color indicates density.

In M.HhaI, the conformational landscape is similar, regardless of apo or bound state and regardless of bound analog. Conversely, in M.TaqI, we can observe two independent free energy wells when we observe the conformational fluctuations of the apo protein as compared to bound with SAH. The apo protein visits conformations in slightly negative values of PC 1, while the SAH-bound protein exhibits fluctuations in the slightly positive values of PC 1. SGH and SUH indicate preference towards negative values of PC 1, or conformations similar to the apo protein. Meanwhile, despite several unbinding events, SCH shows some indication of preference towards a positive PC 1, or a similar conformational landscape to the protein when bound with the SAH analog. Based on experimental results, SAH conformations represent the catalytically competent protein conformations in M.TaqI. In agreement with experimental results which indicate some efficacy with SCM, M.TaqI bound with SCH maintains the catalytically competent conformational space to some proportion during the simulations. Conversely, SGH frequently visits the non-catalytic apo-like conformations, thus elucidating the lack of activity with SGH despite strong binding characteristics.

## Conclusions

Nucleoside bases have a noteworthy role on methylation reaction of methyltransferases. Our results demonstrate that even though the nucleoside base is far from the catalytic centre it dictates the fate of the methylation reaction. Here we prove that M.TaqI specificity for the SAM is governed by holding the adenosine base tightly in the binding pocket and high flexibility by methionine chain. To be specific for SAM, it is important for M.TaqI enzyme to hold nucleoside base tightly and in productive binding mode. While enzyme in engaged in keeping nucleoside tightly it gives more flexibility to methionine chain of SAM where exactly methylation is happening. Unlike M.TaqI, M.HhaI gives more flexibility to the nucleoside base in the binding pocket. Since M.HhaI gives flexibility to nucleoside base, the enzyme is not very careful in choosing the nucleoside base and hence the methylation happens with all SNM analogues. Often adenosine is considered a molecular handle; similarly, our molecular dynamics data indicates that the nucleotide base plays a crucial role in dictating specificity.

Experimentally, we have observed partial methylation by M.TaqI using SCH, but MD simulations showed poor binding for SCH. This data might suggest that tight binding of nucleoside base in the binding pocket is necessary for productive binding hence the methylation. However dynamics of the enzyme also plays an important role as we observed methylation by M.HhaI using all SNH cofactors even after overserved flexibility of the nucleoside base in the binding pocket. To see the effect of similar binding interactions for both enzymes in the binding pocket we made mutant I72W M.TaqI. Experimental data showed that I72W M.TaqI mutant has same activity as M.TaqI (Figure S3). I72W M.TaqI mutant was specific to the SAM. These results suggest that even after the similar interactions in the binding pocket, M.TaqI did not change the specificity. Tryptophane gives more ordered and stronger pi-pi interaction compared to isoleucine, but mutant I72W M.TaqI remains specific as nucleoside base plays key in giving the specificity in case of M.TaqI. Finally, we believe that these novel SNM cofactors might be the good candidates for other classes of methyltransferase (RNA, protein, and small molecule methyltransferase) to explore cofactor engineering as synthetic tool to study methyltransferase role in health and disease.

## Experimental Section

### Materials and methods

ATP, GTP, CTP, UTP, methionine, S adenosylmethionine, Hepes, MgCl^2^, KCl, were purchased from commercial sources and used as supplied unless otherwise mentioned. HhaI methyltransferase, TaqI methyltransferase, TaqI, HhaI, were purchased from New England biolabs. All the experiments were performed using ultrapure water purification system from a MilliQ Integral MT10 type 1 (Millipore).

#### hMAT2A expression and purification

hMAT2A was expressed and purified as reported^[29]^

#### Methylation assay

The standard assay of protection of DNA by methylation against restriction enzyme is used to analyze the methylation. To a reaction mixture containing ATP/GTP/CTP/UTP (5 mM), methionine (10 mM), HEPES (100 mM), MgCl_2_ (10 mM), KCl (50 mM) and hMAT2A (20 μM), pH 8 was incubated for 2 h at 37 °C. After 2 h reaction was quenched by adding cold ethanol, centrifuged for 5 min at 12,000 rpm and the supernatant was transferred to the fresh tube and ethanol was evaporated using speed vac (Genevoc SP scientific miVac modular concentrator series). In the same tube added a 1x cut smart buffer (NEB), DNA (100 ng), M. HhaI (20 U) or M. TaqI (20 U), in a total reaction volume of 20 μL, incubated at 37 °C for M. HhaI and 65 °C for M. TaqI for 1 h. After 1 h enzyme was inactivated by heat for 5 min. Later reaction mixture was charged with 1x cut smart buffer and R. HhaI (10 U) or R. HhaI (10 U) and incubated at 37 °C for HhaI and 65 °C for TaqI for 1 h. The enzyme was inactivated by heat for 5 min and directly loaded onto a 1.5 % agarose gel. The gel was run for 1 h at 100V. Imaged using a ChemiDoc XRS+ gel imaging system (BIORAD).Spontaneous DNA methylation (without DNA methyltransferases), was tested using similar reaction condition for both DNA methyltransferase..

#### Site directed Mutagenesis

Site directed mutagenesis for I72W M.TaqI mutation on M.TaqI plasmid a was carried out using Q5 Site-Directed Mutagenesis Kit (NEB) by following kit protocol and expressed, purified, as reported^[42]^. List of primers is provided in supporting information.

## Acknowledgements

Authors acknowledge to Okinawa Institute of Science and Technology for financial support. This project was supported by a OIST Kick start-up grant. The Laurino lab is supported by a Kakenhi Grant (No. 90812256). We thank Shina Caroline Lynn Kamerlin, Bhanu Pratap Singh Chouhan, Stefano Pascarelli for scientific discussion.

