## Supplementary Information for "Nucleoside-driven specificity of DNA Methyltransferase"

**Substrate for in situ methylation**
The DNA-substrates were prepared from the pet17b plasmid containing tRNA methyltransferases gene using EmeraldAmp PCR Master Mix (Takara Bio Inc.) with forward and reverse primers as mentioned in the table. Synthesized DNA substrate contains the single site of methylation/restriction for TaqI (TCGA), I72W M.TaqI and HhaI (GCGC).

| Name | Forward primer | Reverse primer |
|---|---|---|
| M.TaqI | 5'-AATATAGTTGATGATGGG-3' | 5'-AGGGTGATGGGATGTCC-3' |
| M.HhaI | 5'-AGATCTCGATCCCGCG-3' | 5'-AGAGTCAGACTCTGGGC-3' |

**The DNA-substrate (1810 bp) sequence containing single TaqI restriction or methylation site (TCGA, bold underlined; methylated adenosine in red):**
aatatagttgatgatgggaaatctcagaaacttactcaagatgacataaaagctttgaaggacaagggcattaaaggagaggaaatagttca
gcagttaattgaaaatagtacaacattccgagacaagacagaatttgcccaagataaatatattaaaaagaagaaaaaaaaatatgaagccat
cattactgttgtgaagccatccacccgtattctttcaattatgtattatgcaagagaacctggaaaaattaaccacatgagatacgatacactagc
ccagatgttgacgttgggaaatatccgtgctggcaacaaaatgattgtgatggaaacgtgtgcaggcttggtgctgggtgcaatgatggaac
gaatgggaggttttggctccattattcagctataccctggaggaggacctgttcgggcagcaacagcatgttttggatttcccaaatcttttctca
gtggtctttatgaattccctctcaacaaagtggacagtcttctacatggaacattttctgccaagatgttatcttcagagccaaaagacagtgcttt
ggttgaagaaagtaatggcacactggaggaaaaacaggcttctgaacaagagaatgaagacagcatggcagaggccccagagagcaac
cacccagaagaccaggaaacaatggaaacaatttctcaagatccagaacataaggggcctaaagagagaggaagcaaaaaagattatatt
caggaaaaacagaggagacaagaagagcagaggaaaagacatttagaggctgccgctctgctgagtgaaagaaacgcagatggtttaat
tgtagctagtcgtttccaccccactcccctgctgctgtctttgctggactttgtggccccttcaaggccgtttgtggtctactgtcagtacaaaga
gcctctgttggaatgctacacaaaactgcgggagaggggaggggtcatcaacctcaggctgtctgaaacctggctcagaaattatcaggttt
tgccaga**TCGA**agtcatcctaaactgctgatgagtggaggtgggggttatcttctctccggcttcaccgttgccatggacaaccttaaagc
agacaccagcctcaaatctaatgcaagcactttagaatcacacgagactgaggagcctgcagctaaaaaacgaaaatgcccagagtctga
ctcttaagcatctagaaataattttgtttaactttaagaaggagatatacatatgagcttcgtggcatacgaggagctgatcaaggagggtgaca
cggccatcctgtcactgggccatggtgcaatggtggcagtgcgtgtgcagcgtggggcacagacccagacccggcatggtgtcctgcgg
cactcagttgaccttatcggccgccccttcggctccaaggtgacgtgcggccgaggtggctgggtgtatgtgctgcaccccacgcccgag
ctctggacgctgaacctgccgcaccgcacgcagatcctctactccacagacatcgccctcatcaccatgatgttggagcttcggcccggctc
tgtggtctgtgagtctggcaccggcagtggctctgtgtcccacgccatcatccgcaccattgcacccacgggtcacctgcacacggtggag

ttccaccagcagcgggcagagaaggcccgggaggagttccaggagcaccgtgtgggccgctgggtgactgtgcgcacccaggacgtg
tgccgcagtggctttggcgtgagccacgtggccgacgccgtcttcctggacatcccatcaccct

**The DNA-substrate (1723bp) sequence containing single HhaI restriction or methylation site (GCGC, bold underlined; methylated cytosine in red):**

agatctcgatcccgcgaaattaatacgactcactatagggagaccacaacggtttccctctagaaataattttgtttaactttaagaaggagata
tacatatggctggttgcctgaacgacatcttcgaagctcagaaaatcgaatggcaccatcaccatcaccatctggttccgcgtggttccggcg
gcggcggcctgcagcttgaagtcctctttcagggacccg**GCGC**catggagggctcaggggagcagccgggcccacaaccacagca
tcccggagaccaccgcatccgcgacggcgacttcgtggtgctgaaacgtgaagatgtgtttaaagcagtacaagtccagcggagaaaaaa
agtaactttcgaaaaacagtggttctacctggataacgtcattggccatagttatggaactgcatttgaagtgaccagtggaggaagtctacag
cccaagaagaagagggaagagcctactgcagagactaaagaagcgggcactgataatcgaaatatagttgatgatgggaaatctcagaa
acttactcaagatgacataaaagctttgaaggacaagggcattaaaggagaggaaatagttcagcagttaattgaaaatagtacaacattccg
agacaagacagaatttgcccaagataaatatattaaaaagaagaaaaaaaaaatatgaagccatcattactgttgtgaagccatccacccgtat
tctttcaattatgtattatgcaagagaacctggaaaaattaaccacatgagatacgatacactagcccagatgttgacgttgggaaatatccgtg
ctggcaacaaaatgattgtgatggaaacgtgtgcaggcttggtgctgggtgcaatgatggaacgaatgggaggttttggctccattattcagc
tataccctggaggaggacctgttcgggcagcaacagcatgttttggatttcccaaatcttttctcagtggtctttatgaattccctctcaacaaag
tggacagtcttctacatggaacattttctgccaagatgttatcttcagagccaaaagacagtgctttggttgaagaaagtaatggcacactgga
ggaaaaacaggcttctgaacaagagaatgaagacagcatggcagaggccccagagagcaaccacccagaagaccaggaaacaatgga
aacaatttctcaagatccagaacataaggggcctaaagagagaggaagcaaaaaagattatattcaggaaaaacagaggagacaagaag
agcagaggaaaagacatttagaggctgccgctctgctgagtgaaagaaacgcagatggtttaattgtagctagtcgtttccaccccactccc
ctgctgctgtctttgctggactttgtggccccttcaaggccgtttgtggtctactgtcagtacaaagagcctctgttggaatgctacacaaaactg
cgggagaggggaggggtcatcaacctcaggctgtctgaaacctggctcagaaattatcaggttttgccagatcgaagtcatcctaaactgct
gatgagtggaggtgggggttatcttctctccggcttcaccgttgccatggacaaccttaaagcagacaccagcctcaaatctaatgcaagca
ctttagaatcacacgagactgaggagcctgcagctaaaaaacgaaaatgcccagagtctgactct

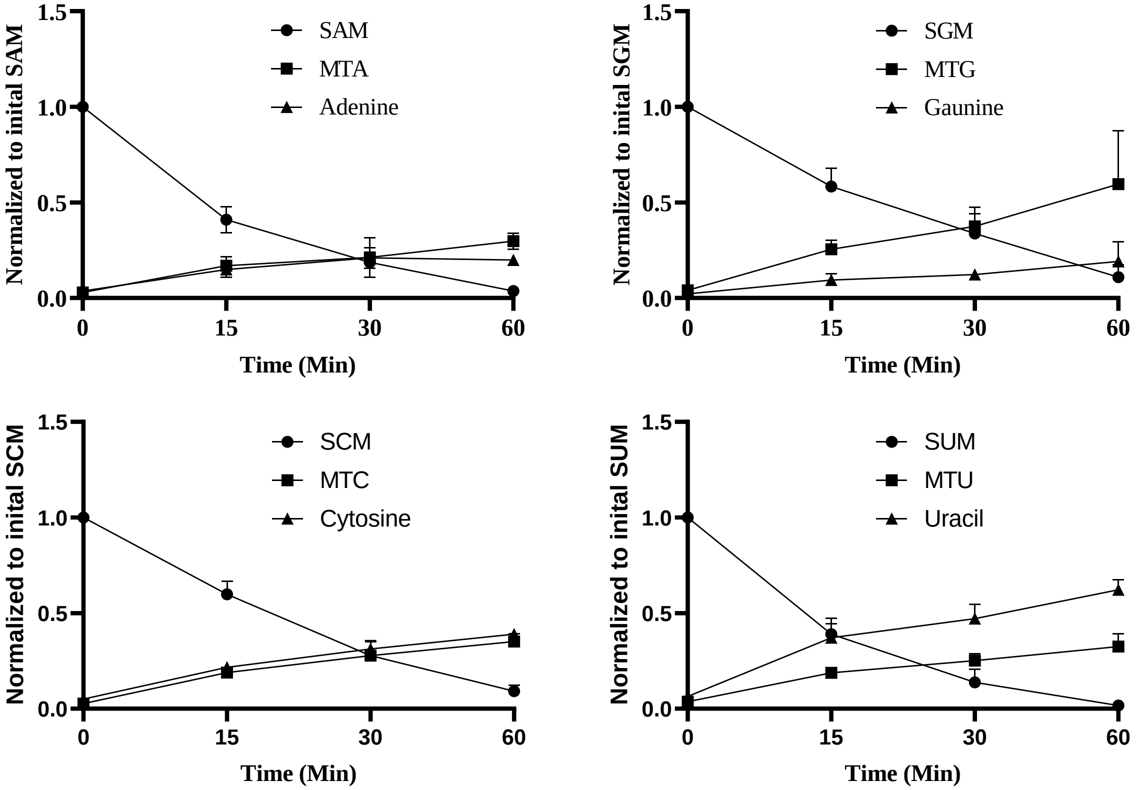


**Figure S1: SNM stability at 65 °C**. SNM analogs were incubated at 65 ℃ and aliquots were taken after 15, 30 and 60 minutes. Stability of each sample was tested by UPLC.[1] SNM analog degrades to the corresponding MTN and nucleotide bases with respect to the time. Normalized graph with corresponding SNM. Experiments were run in duplicate and error bars shows standard deviation.

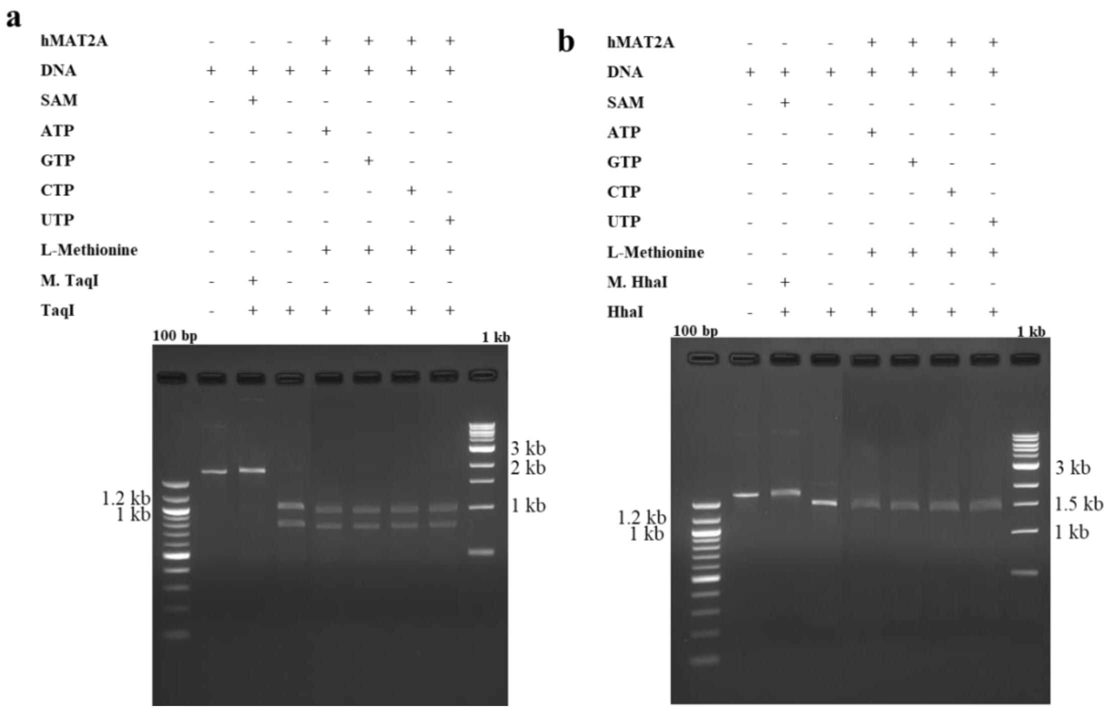


**Figure S2: Methylation without DNA methyltransferases enzyme** a) M. TaqI. methylation without enzyme b) M.HhaI methylation without enzyme, using 1.5 % agarose gel at 100 V for 1hr 1 kb and 100 bp DNA ladder (NEB).

### Site directed Mutagenesis

The primers used for mutagenesis listed below.

| Name | Forward primer | Reverse primer |
| --- | --- | --- |
| I72W M.TaqI | 5'- TGGGGGTAGAGTGGGACCCTAAGGCG-3' | 5'- CGAAGCGGTATGCAGTCCCATGTG -3' |

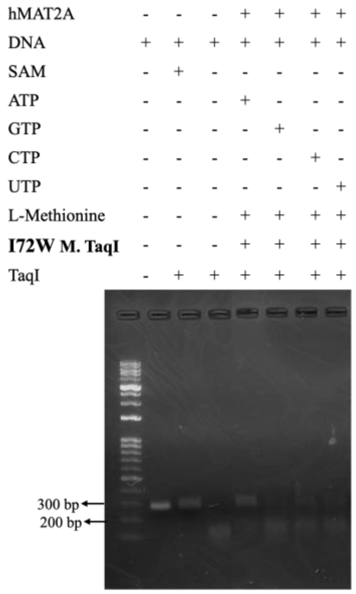


**Figure S3: Agarose gel analysis of DNA modification** (a) Enzymatic modification by I72W M.TaqI using SNM cofactors. Agarose gel was 1.5 % run for 1 hr. at 100 V, stained with ethidium bromide. GeneRuler DNA ladder 10kb was used.
